# The protector within: Comparative genomics of APSE phages across aphids reveals rampant recombination and diverse toxin arsenals

**DOI:** 10.1101/2020.03.17.996009

**Authors:** Jeff Rouïl, Emmanuelle Jousselin, Armelle Coeur d’acier, Corinne Cruaud, Alejandro Manzano-Marín

## Abstract

Phages can fundamentally alter the physiology and metabolism of their hosts. While ubiquitous in the bacterial world, they have seldom been described among endosymbiotic bacteria. One notable exception in the APSE phage that is found associated with the gammaproteobacterial *Hamiltonella defensa*, hosted by several insect species. This secondary facultative endosymbiont is not necessary for the survival of its hosts but can infect certain individuals or even whole populations. Its infection in aphids is often associated with protection against parasitoid wasps. This protective phenotype has actually been linked to the infection of the symbiont strain with an APSE, which carries a toxin cassette that varies among so-called ”types”. In the present work, we seek to expand our understanding of the diversity of APSE phages as well as the relations of their *Hamiltonella* hosts. For this, we assembled and annotated the full genomes of 16 APSE phages infecting *Hamiltonella* symbionts across 10 insect species. Molecular and phylogenetic analyses suggest that recombination has occurred repeatedly among lineages. Comparative genomics of the phage genomes revealed two variable regions that are useful for phage typing. Additionally, we find that mobile elements could play a role in the acquisition of new genes in the toxin cassette. Altogether, we provide an unprecedented view of APSE diversity and their genome evolution across aphids. This genomic investigation will provide a valuable resource for the design and interpretation of experiments aiming at understanding the protective phenotype these phages confer to their insect hosts.

## Introduction

Bacteriophages, the viruses that infect and replicate in bacteria, exhibit genomes that are highly dynamic (reviewed in Dion *et al.*, 2020). They undergo rampant recombination and constant rearrangements. They are also known to play a prominent role in horizontal gene transfers between bacteria (reviewed in Touchon *et al.*, 2017). As such, they can be a source of innovation for the bacteria that are infected as well as for the eukaryotic organisms that carry these bacteria. Aphids (Hemiptera: Aphididae) can host a wide variety of facultative endosymbiotic bacteria (Guo *et al.*, 2017), in addition to their obligate nutritional endosymbiont *Buchnera aphidicola*. These bacteria are not necessary for reproduction nor survival, but can endow their host with a variety of beneficial effects, ranging from survival after heat stress to defence against pathogenic fungi and parasitoid wasps (reviewed in Oliver *et al.* 2010). Although bacteriophages are ubiquitous wherever bacteria exist, their discovery is scarce among the endosymbiotic bacteria of insects. One such case is the APSE phage: a lambdoid phage with an isometric head and a short tail (resembling species in the family *Podoviridae*) which was originally isolated from an *Acyrthosiphon pisum* aphid infected by an unidentified secondary endosymbiont (van der Wilk *et al.*, 1999). This phage has been found so far associated with the gammaproteobacterial symbiont species *Hamiltonella defensa*: an endosymbiont lineage that infects several hemipteran species. In the case of aphids, it is clear that toxin genes carried by this phage confer a protective phenotype against parasitoid wasps by disabling wasp development (Brandt *et al.*, 2017; Oliver *et al.*, 2009).

Microscopic analysis and genome sequencing of this phage, first termed APSE-1 (for bacteriophage **1** from ***Ac***. ***p**isum* **s**econdary **e**ndosymbiont), revealed a circularly permuted and terminally redundant double-stranded DNA molecule of 36,524 base pairs (**bp**) (van der Wilk *et al.*, 1999). APSE-1 encodes for a protein product showing low sequence similarity with the shiga-like toxin B subunit of several bacteriophages. Further whole genome sequencing of a second APSE genome from *Ac. pisum* (termed APSE-2) as well as specific regions of other APSE phages revealed variation across the so-called ”types” (Degnan and Moran, 2008; Moran *et al.*, 2005). These ”types” are characterised by carrying a unique set of genes including (putative) toxins from three protein families: Shiga-like toxin, cytolethal distending toxin (**CdtB**), and YD-repeat toxin. So far seven types of APSE have been described (table 1). They are associated with *Hamiltonella* strains from six aphid species (from two subfamilies) and the whitefly *Bemisia tabaci*.

**Table 1.** Previously characterised APSE toxins. Summary of the previous works characterising APSE phage toxins. + Later referred to as APSE-8 (Brandt *et al.*, 2017). * Variants of APSE-1, see Degnan and Moran (2008). NA= Not available.

| Insect host | *Hamiltonella* strain | APSE type | Toxin | Reference |
| --- | --- | --- | --- | --- |
| *Ac. pisum* | NA | APSE-1 | Shigga-like | van der Wilk *et al.* (1999) |
| *Ac. pisum* | 5AT | APSE-2 | CdtB | Degnan and Moran (2008); Moran *et al.* (2005) |
| *Ac. pisum* | NY26 | APSE-2 | CdtB | Brandt *et al.* (2017) |
| *Ac. pisum* | 82B | APSE-2 | CdtB | Degnan and Moran (2008); Moran *et al.* (2005) |
| *Ac. pisum* | ZA17 | APSE-2<sup>+</sup> | CdtB | Martinez *et al.* (2014) |
| *Ac. pisum* | WA4 | APSE-2 | CdtB | Martinez *et al.* (2014) |
| *Ac. pisum* | A1A,A2F,A2H | APSE-3 | YD-repeat | Degnan and Moran (2008) |
| *Ac. pisum* | AS3 | APSE-3 | YD-repeat | Oliver *et al.* (2009) |
| *Ac. pisum* | AS5 | APSE-3 | YD-repeat | Oliver *et al.* (2009) |
| *Ac. pisum* | R7 | APSE-3 | YD-repeat | Oliver *et al.* (2009) |
| *Aphis fabae* | H76 | APSE-3 | YD-repeat | Dennis *et al.* (2017) |
| *Aphis craccivora* | 5ATac | APSE-4* | Shigga-like | Degnan and Moran (2008) |
| *Uroleucon rudbeckiae* | NA | APSE-5* | Shigga-like | Degnan and Moran (2008) |
| *Chaitophorus* sp. | N4 | APSE-6 | CdtB | Degnan and Moran (2008) |
| *Aphis fabae* | H402 | APSE-6 | CdtB | Dennis *et al.* (2017) |
| *Bemisia tabaci* | NA | APSE-7 | CdtB | Degnan and Moran (2008) |

Differences in the level of protection have been observed in different associations of aphids species/genotypes and *Hamiltonella*-APSE combinations. For example, experimental work has shown that *Hamiltonella* endosymbionts can confer varying levels of resistance against different parasitoid wasp species, and even no defence at all (Asplen *et al.*, 2014; Hopper *et al.*, 2018; Lenhart and White, 2017; Leybourne *et al.*, 2020). It has been shown that *Hamiltonella* carrying APSE-3 confer a strong defensive phenotype, while the ones carrying APSE-2 generally confer moderate defence against parasitoids (Brandt *et al.*, 2017; Martinez *et al.*, 2014; Oliver *et al.*, 2009). This protection is indeed largely dependant on the APSE ”type” (Degnan and Moran, 2008; Martinez *et al.*, 2014), and experiments have conclusively demonstrated that APSE-3, carried by *Hamiltonella* strain AS3, can confer its protective phenotype when transplanted to the naturally-occurring APSE-free and non-protective strain A2C (Brandt *et al.*, 2017). The beneficial fitness effect of APSE-bearing *Hamiltonella* has been shown to be conditional to the presence of APSE and the environmental pressure of parasitoid infection. In one study, the authors found that, while apparently no cost to infection with *Hamiltonella* could be detected in aphids not exposed to parasitism, a significant decline in the frequency of *Hamiltonella* was observed (Oliver *et al.*, 2008). In another study, the loss of APSE led to an increase in intra-aphid *Hamiltonella* abundance and was associated to ”severe” deleterious effects on aphid fitness (Weldon *et al.*, 2013).

In the current work we sought to explore and expand our knowledge on the diversity of APSE phages across aphid species. For this purpose, we assembled and annotated full genomes for 16 APSE phages that infect *Hamiltonella* endosymbionts across 10 insect species, including several aphids (from 5 subfamilies) and the whitefly *Bemisia tabaci*. We performed phylogenetic analyses on *Hamiltonella* and its associated APSE phages in order to understand the diversity and evolutionary trajectory of this defensive phage. Through comparative genomics, we investigated genome rearrangement and more specifically look at the evolution of the toxin cassette. We found evidence suggesting that recombination takes place among APSE types in aphids and that most variation in gene content is observed in two main regions of their genomes. Finally, analysis of the toxin cassettes revealed that mobile elements might be involved in at least some of the variation observed in this symbiotically relevant genomic region.

## Materials and Methods

### Aphid collection, DNA extraction, and sequencing

Complete genome sequences of APSE phages were retrieved from the NCBI database for APSE-1 (also known as *Hamiltonella virus APSE1*) and *Hamiltonella* strain 5AT. We then used the genome data available for five *Hamiltonella* strains from the aphid *Ac. pisum* and two from the whitefly *B. tabaci* to extract integrated APSE phages. Additionally, we gathered sequencing data from eight different aphid species (from four genera belonging to four subfamilies) collected between the years 2008 and 2016: some of which were previously known to host *Hamiltonella* (Meseguer *et al.*, 2017). Specimens were kept in 70% ethanol at 6°C. For whole-genome sequencing, we prepared DNA samples enriched with bacteria following a slightly modified version of the protocol by Charles and Ishikawa (1999) as described in Jousselin *et al.* (2016). For this filtration protocol *circa* 15 aphids from one colony were pooled together. Extracted DNA was used to prepare custom paired-end libraries in Genoscope as in Manzano-Marín *et al.* (2018). These libraries were sequenced using either 151 or 251 bp paired-end reads chemistry on a HiSeq2500 Illumina sequencer. For full details on specimen collection, species identification, and accession numbers for the samples from which APSE phages were extracted, see supplementary table S1.

### *Hamiltonella* and APSE genome assembly and annotation

Firstly, we scanned the afore-mentioned *Hamiltonella* assembled genomes using the sequence of the phage attachment site (also known as **attP**) and the last 62 bp with a 90% identity threshold in **UGENE** v1.29.0 (Okonechnikov *et al.*, 2012). For non-circularised genomes, we first scanned the contigs using blastn (Altschul, 1997) for these sequences in order to identify the contig or scaffold where the putative APSE phage resided in. Illumina sequences from the eight newly sampled aphid species were first right-tail clipped (requiring a minimum quality threshold of 20 and a minimum length of 75 bp) using **FASTX-Toolkit** v0.0.14 (http://hannonlab.cshl.edu/fastxtoolkit/, last accessed, March 4 2020). Additionally, **PRINSEQ** v0.20.4 (Schmieder and Edwards, 2011) was used to remove reads containing undefined nucleotides as well as those left without a pair after the filtering and clipping process. The resulting reads were assembled using **SPAdes** v3.11.1 (Bankevich *et al.*, 2012) with the --only-assembler option and k-mer sizes of 33, 55, 77, 99, and 127. From the resulting contigs, those that were shorter than 200 bps were dropped. The remaining contigs were binned using results from a **BLASTX** (Altschul, 1997) search (best hit per contig) against a database consisting of the Pea aphid’s proteome and a selection of aphid’s symbiotic bacteria proteomes (supplementary table S2). When no genome was available for a specific bacterial lineage, closely related bacteria were used. The assigned contigs were manually screened using the **BLASTX** web server (searching against the nr database) to ensure correct assignment. This binning process confirmed the presence of *Buchnera*, *Hamiltonella*, and its corresponding APSE phage. APSE was confidently assigned to *Hamiltonella* by checking for chimeric APSE-*Hamiltonella* chromosome contig ends in the corresponding insertion sites and/or by checking that no other facultative symbionts were sequenced in the sample. The resulting contigs were then used as reference for read mapping and individual genome assembly using **SPAdes**, as described above. *Hamiltonella* draft assemblies included APSE-assigned reads. APSE individual assemblies were visually screened for inconsistencies using Tablet (Milne *et al.*, 2013).

The resulting APSE genomes underwent a draft annotation using **Prokka** 1.14.4 (Seemann, 2014). This was followed by non-coding RNA prediction using **infernal** v1.1.2 (Nawrocki and Eddy, 2013) (against the **Rfam** v14.1 database (Kalvari *et al.*, 2018a, b)) and **tRNAscan-SE** v2.0.5 (Chan *et al.*, 2019). We then performed manual curation of the annotations on **UGENE** (Okonechnikov *et al.*, 2012) through on-line **BLASTX** searches of the intergenic regions, open reading frame (**ORF**) finding feature in **UGENE**, and through **DELTA-BLAST** (Boratyn *et al.*, 2012) searches of the predicted ORFs against NCBI’s nr database and the **InterProScan** v5 web server (Jones *et al.*, 2014; Mitchell *et al.*, 2019). **SignalP** v5.0 (Almagro Armenteros *et al.*, 2019) and **Phobius** v1.01 (Kall *et al.*, 2005) were used to predict signal peptides. ORFs were considered to be putative functional proteins (and thus not pseudogenes) if seemingly essential domains for the function were found or if the ORFs displayed truncations but retained identifiable domains. Short ORFs (≤300) that were conserved but consistently pseudogenised in all but one APSE genome were annotated as miscellaneous features. The origin of replication was determined using **originX** (Worning *et al.*, 2006). Once the first APSE genome was curated, we used these proteins as a ”genus”-specific database in **Prokka** to annotate the subsequent genomes and iteratively added novel proteins to the database. Codon usage was calculated separately for closed high-quality *Hamiltonella*-APSE pairs using the **Cousin** web sever (Bourret *et al.*, 2019).

Finally, phage-like regions were searched in *Arsenophonus* spp. genomes using the web server for **PHASTER** (Arndt *et al.*, 2016; Zhou *et al.*, 2011). The retrieved phage-like regions were then manually inspected in **UGENE** and identity of genes *vs.* those of APSE was assessed using the online web server of **BLAST**. The **EMBOSS** suite program **primersearch** (Rice *et al.*, 2000) was used to predict possible amplification targets in the *Arsenophonus* genomes using the primers reported in Duron (2014).

### Phylogenetic and recombination analyses

In order to reconstruct the phylogeny of *Hamiltonella* endosymbionts, we used seven gene sequences (*accD*, *dnaA*, *gyrB*, *hrpA*, *murE*, *ptsI*, and *recJ*) following Degnan and Moran (2008). Genes sequences were gathered from the Supplementary data for Manzano-Marín *et al.* (2020) (https://doi.org/10.5281/zenodo.2566355, last accessed March 4, 2020) then used to identify orthologs in newly sequenced *Hamiltonella* strains using **BLASTX**. Due to the low coverage for *Hamiltonella* draft genomes for two samples (3702 and 3692) we were unable to recover sequences for all seven genes used for phylogenetic inference, and thus these specimens were excluded from the *Hamiltonella* phylogenetic analysis. For each gene, sequence alignments were conducted using **MUSCLE** v.3.8.31 (Edgar, 2004) and visually checked using **SeaView** v4 (Gouy *et al.*, 2010). We then removed divergent and ambiguously aligned blocks using **Gblocks** v0.91b (Talavera and Castresana, 2007). A partitioned scheme was selected using **PartitionFinder** v2.1.1 (Lanfear *et al.*, 2017) with one data block defined for each codon position (1^st^, 2^nd^, and 3^rd^) in each gene. The best model of evolution of each partition was selected among those available in **MrBayes**, using the Bayesian Information Criterion metric under a *greedy* algorithm. Finally, we concatenated the resulting alignments and ran a Bayesian phylogenetic inference using the GTR+I+G model in **MrBayes** v3.2.7 (Ronquist *et al.*, 2012) running two independent analyses with four chains each for 1,500,000 generations and checked for convergence.

For performing both phylogenetic inferences and analysing the genetic differences across APSE, we first ran an orthologous protein clustering analysis using **OrthoMCL** v2.0.9 (Chen *et al.*, 2007; Li, 2003) on the full predicted proteome for the phage genomes (supplementary table S3). We then extracted the single copy-core proteins for phylogenetic reconstruction (30 protein groups). Sequences were aligned using **MAFFT** v7.450 (maxiterate 1000 localpair) (Katoh and Standley, 2013). Divergent and ambiguously aligned blocks were removed using **Gblocks**. Substitution model was selected using **ModelTest-NG** v0.1.6 (Darriba *et al.*, 2020) by the AIC criterium. Bayesian phylogenetic reconstruction was conducted with **MrBayes** as described above by running chains for 300,000 generations and checking for convergence. Given prior evidence for intragenic recombination in APSE proteins (Degnan and Moran, 2007), we constructed a second dataset removing all genes where recombination has putatively occurred. Recombination was tested for running **PhiPack** (Bruen *et al.*, 2006) on the aligned proteins and results of these analyses, for the putative non-recombining proteins based on Φ_*ω*_, can be found in supplementary table S4. Bayesian phylogenetic reconstruction was ran as described above.

To infer the relationships of the 14 toxin-cassette- and lytic-region-proteins (lysozyme and holins) in APSE genomes, we searched for similar sequences using the online BLASTP web server *vs.* the NCBI’s nr database and selected the top 50 non-redundant hits. These were filtered for a minimum of 70% query coverage and a 35% sequence identity. We aligned each of them using **MAFFT** and manually removed divergent and ambiguously aligned blocks. For each gene, model selection was done with **ModelTest-NG**. Bayesian phylogenetic inference was run for 300,000 generations in **MrBayes** as described above.

All resulting trees were visualized and exported with **FigTree** v1.4.1 (http://tree.bio.ed.ac.uk/software/figtree/, last accessed March 4, 2020). All files used for phylogenetic analyses as well as **PHASTER** phage search results and comparative genomics files used in this study are available in https://doi.org/10.5281/zenodo.3764739 (last accessed April 24, 2020).

## Results and Discussion

### APSE phage genomes

We successfully *de novo* assembled genomes for eight complete APSE phages from eight different aphid species and extracted an additional eight from both closed and draft *Hamiltonella* endosymbionts from the pea aphid *Ac. pisum* and the whitefly *B. tabaci* (table 2). Additionally, we recovered draft genomes for all eight *Hamiltonella* strains hosting the newly sequenced APSE phages. All recovered phage genomes have a genome size between 33,476 and 41,343 bp. They have a very conserved G+C content ranging between 42.81% to 45.04% and code for an average of 38 CDSs with 3 to no pseudogenes (table 2). Most APSE phages preserve an intact tRNA-Lys-(UC)UU, which has been lost in 5D and ZA17, both of which still keep an internally-degraded tRNA pseudogene. An analysis of 17 bacterial hosts and 37 phage genomes found that the presence of tRNAs in these phages tended to correspond to codons that were highly used by the phage’s protein coding genes but were rare in the host’s (Bailly-Bechet *et al.*, 2007). However, we found virtually no difference in the lysin codon usage between pairs of *Hamiltonella*-APSE genomes (supplementary table S5). Nonetheless, we found that the *Hamiltonella* strain 5AT only codes for a tRNA-Lys-CUU, following the pseudogenisation of the conserved tRNA-Lys-UUU, which recognises the most commonly used codon for Lysine. Conversely, its associated APSE preserves its own tRNA-Lys-UUU. This suggests that the presence of a tRNA-Lys-UUU in the APSE phage could drive the loss of this tRNA in the bacterial host, which could in turn provide a selective advantage to carrying this phage.

**Table 2.**
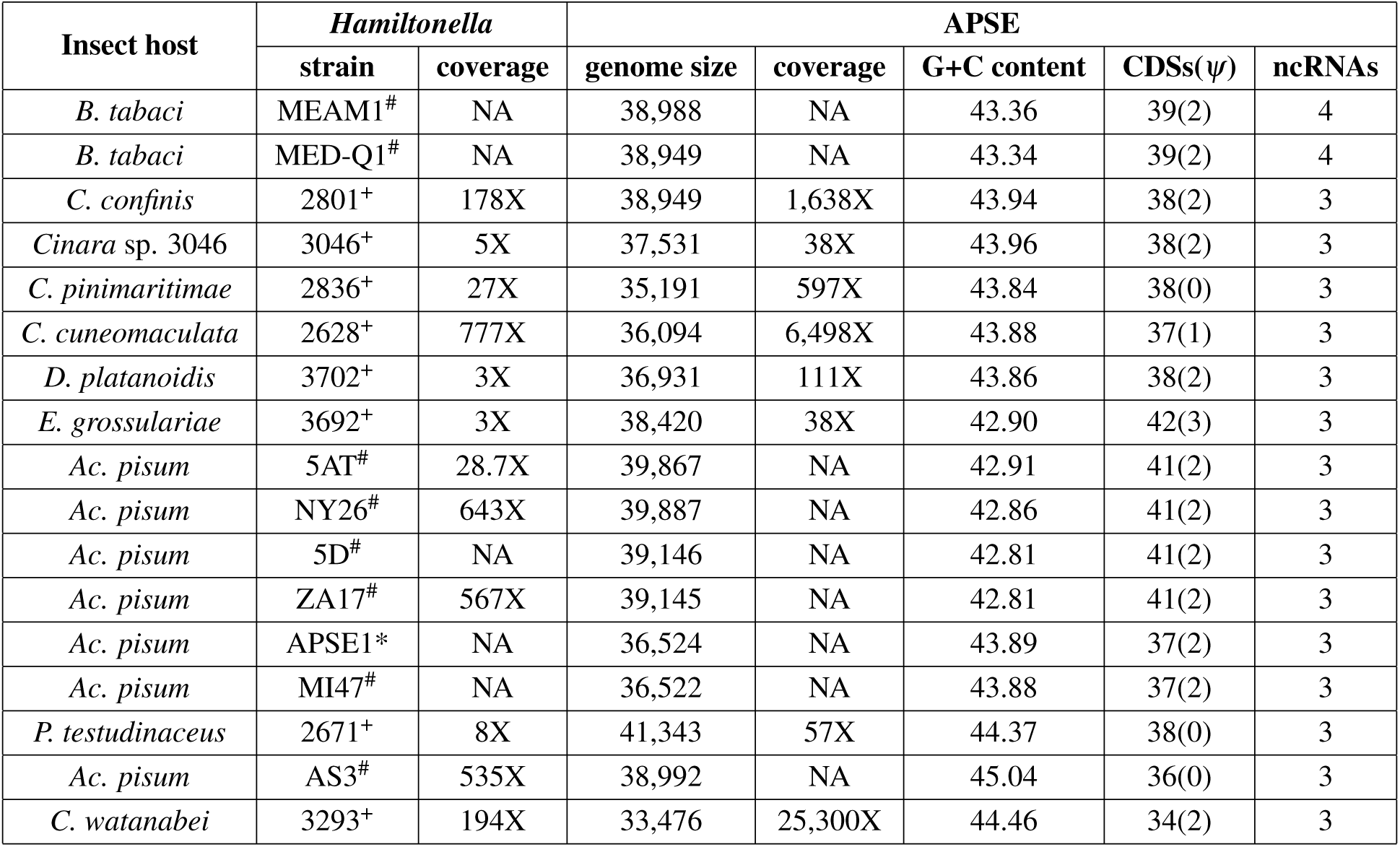
Genome assembly and annotation statistics. Assembly and genome statistics for complete APSE genomes and their *Hamiltonella* hosts. # Extracted from publicly available *Hamiltonella* genomes. + Sequenced in this study. * Phage isolated from an unidentified secondary symbiont of *Ac. pisum* (van der Wilk *et al.*, 1999). NA= Not available.

All APSE have two non-coding RNAs (**ncRNAs**) that code for small bacterial RNAs (**sRNA**): a c4 antisense family sRNA and an isrK Hfq binding family sRNA. In the case of the former, a c4 sRNA has been identified in P1 and P7 phages from *Escherichia coli* as a regulator of the *ant* gene, an anti-repressor, through the binding to complementary regions in the upstream region of this gene (Citron and Schuster, 1990). Through the analysis of the putative so-called a’ and b’ antisense regions of the c4 sRNA, we found that they consistently matched the 3’-end region of the distant DNA polymerase gene and the upstream non-coding region of the adjacent BRO family, N-terminal domain-contaning protein. Three BRO family proteins (BRO-A, BRO-C, and BRO-D) from the *Bombyx mori nucleopolyhedrovirus* (BmNPV) have been suggested to influence host DNA replication and/or transcription, following the finding that they are able to bind DNA from the host with a stronger affinity for single-stranded than for double-stranded DNA (Zemskov *et al.*, 2000). Regarding IsrK, its expression pattern in *Salmonella typhimurium* suggested that this sRNA might be involved in the regulation of virulence (Padalon-Brauch *et al.*, 2008). Lastly, both APSE phages harboured by *Hamiltonella* infecting MEAM1 and MED-Q1 *B. tabaci* biotypes code for an additional sRNA-Xcc1 family sRNA upstream of a protein of unknown function (ORF6N domain-containing protein) also unique to these APSE. In *Xanthomonas campestris* pv. *vesicatoria*, *sRNA-Xcc1* is under the positive control of two important virulence regulators, suggesting that they might play a role in pathogenesis (Chen *et al.*, 2011). All these points towards an important regulatory function of these sRNAs in APSE and highlights them as targets to study their downstream regulatory effects in both APSE and *Hamiltonella* gene expression.

### APSE gene content variation and toxin cassettes

APSE genomes were highly conserved in both gene content, order, and sequence identity (supplementary fig. S1, table S3, and file S3). The most variable genes, in terms of sequence identity, across APSE were those of a BRO family, N-terminal domain-containing protein (orthogroup APSEcp_004); the DNA polymerase (APSEcp_008); the lyzozyme (APSElp 003); the small subunit of the terminase (APSEcp_020); and the tail proteins (needle, spike, and fiber assembly; APSEcp_028 and APSEcp_033-34). Regarding the DNA polymerase gene, we observed differential loss of the intein domain (a protein intron). The lyzozyme gene was by far the one that showed the most sequence divergence. A phylogenetic analysis suggested two well supported lyzozyme phylogroups (so-called ”P13” and ”F”, supplementary fig. S2), consistent with previous findings (Degnan and Moran, 2008). From our analyses, ”P13” is restricted to APSE-7 and APSE-1 (including the APSE-1 variants APSE-4 and APSE-5).

One marked difference in CDS content between APSE harboured by *Hamiltonella* symbionts of aphids and those harboured by the whitefly *B. tabaci* was in the protein flanked by the BRO family, N-terminal domain-containing protein and the putative P-loop NTPase. While all APSE from aphid-infecting *Hamiltonella* code for a putative phage regulatory protein (Rha family), those infecting the whitefly code for an AntA/AntB antirepressor family protein. To our knowledge, the function of both of these proteins is not well understood, and thus the significance of this difference remains to be studied.

Altogether, we identified two main regions of genomic variation in APSE phages, and propose an APSE classification based on the composition of these two variable sequence stretches based on Degnan and Moran (2008), and keeping the main type numbering (X), with a slight revision (X.Yz) based on our findings (fig. 1): The first variable region (X) is the toxin cassette, with small variations within the types (Y), and the second one is found around the vicinity of the DNA polymerase gene (z). In regards to the toxin cassettes, we found five different ”types” or ”gene sets” represented in the newly assembled APSE genomes. We found seven phages that could be classified within APSE-7. These were associated with *Hamiltonella* from two whiteflies and five species of aphids (from two subfamilies). We found that while the CdtB family protein was conserved across APSE-7, its companion hypothetical protein was not. Both proteins in the cassette showed identifiable signal peptides, suggesting their export. The CdtB family protein is also present in APSE-2, however the structure of the cassette is unlike that of APSE-7. It codes for an additional two putative toxin genes, and in the case of APSE-2.2 subtypes, two flanking hypothetical proteins, all with signal peptides. In the case of APSE-2.1a, we found that the hypothetical proteins flanking CdtB were missing. Nonetheless, we found a different hypothetical protein (containing a signal peptide) flanked by insertion sequence pseudogenes. This suggests that, at least in this case, the new member of the toxin cassette was likely mobilised by the action of transposable elements. As in a previous study by (Degnan and Moran, 2008), we found that all APSE-2 and −7 types had both holin family genes (lambda and HP1) followed by a different lysozyme gene phylotype, so-called ”F” and ”P13” respectively. The APSE-1 type was represented by two nearly identical APSE genomes, both associated with *Hamiltonella* strains from *Ac. pisum*. Both presented the same four hypothetical proteins, including two previously identified as putative subunits of a shiga-like toxin (Degnan and Moran, 2008; van der Wilk *et al.*, 1999). APSE-3 types were found to be associated with *Hamiltonella* hosted by two distantly related aphid species, *Ac. pisum* and *P. testudinaceus*. They showed a very conserved cassette, encoding for a hypothetical protein, with no identifiable signal peptide, and an RHS-repeat putative toxin (previously referred to as ”YD-repeat-containing” by Degnan and Moran 2008) with a signal peptide. Finally, the cassette found in 3293 (*Hamiltonella* from *C. watanabei*) represents a completely novel type, which we classified as APSE-8. This toxin cassette encodes for a single putative virulence-associated protein with no recognisable signal peptide. Previous studies have classified the APSE-2 variant found in *Hamiltonella* strain ZA17 as APSE-8 (Brandt *et al.*, 2017; Chevignon *et al.*, 2018; Doremus and Oliver, 2017; Patel *et al.*, 2019). However, we found that the only difference in its toxin cassette was the pseudogenisation of the first putative toxin gene within this region when compared to APSE-2 5AT and NY26. Therefore, we have considered it to be a subtype of APSE-2, in agreement with Martinez *et al.* (2014).

**Figure 1.**
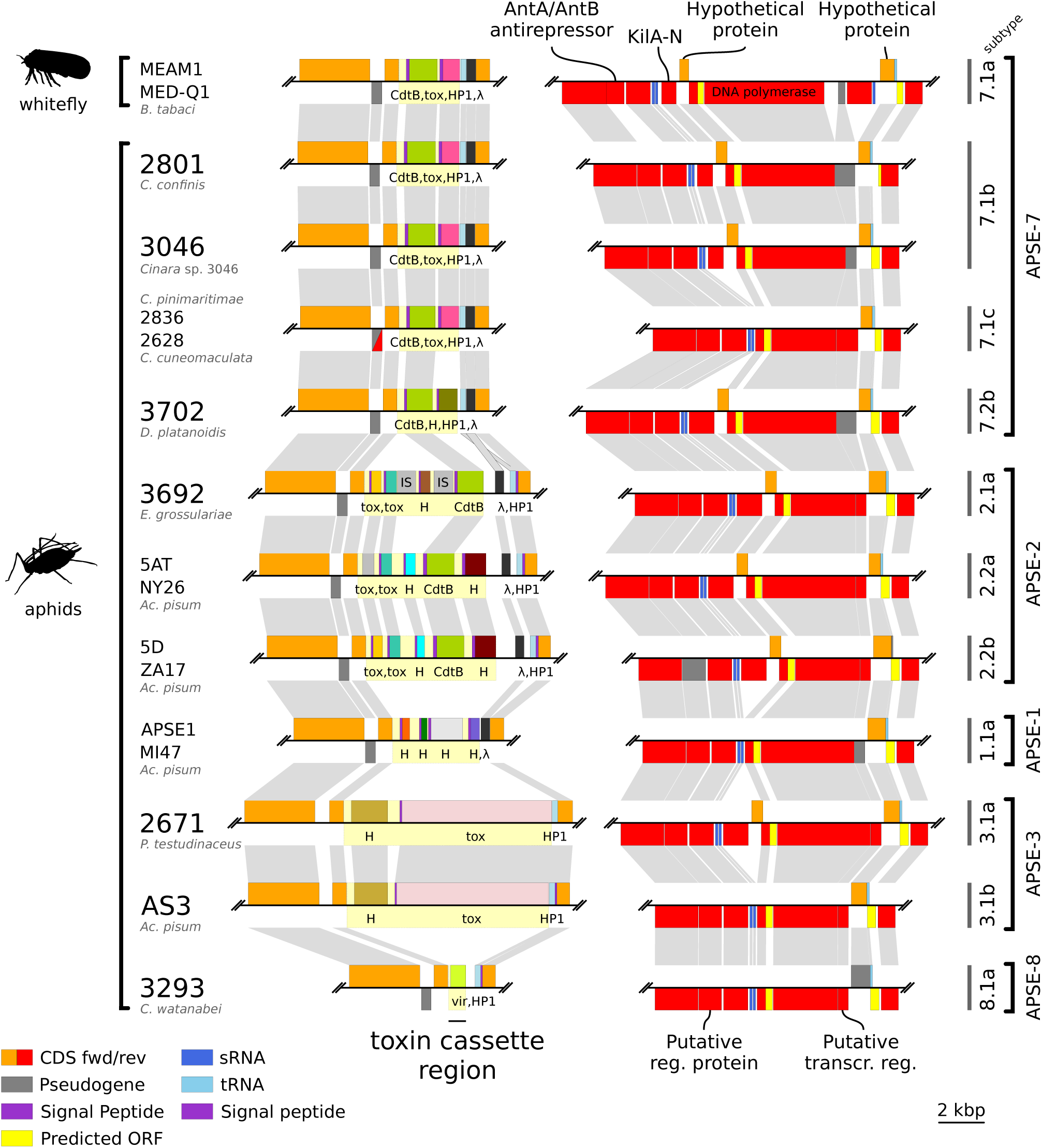
Variable regions in APSE genomes. Genome synteny plots of the two variable regions across APSE phages. On the bottom left, a colour key for the features shown in the plot. On the right, proposed classification of APSE in types and subtypes based on the variable regions composition.

Regarding the second variable region, the main differences laid in the presence/absence of the KilA-N domain-containing protein and its companion hypothetical protein, a putative transcriptional regulator upstream of the DNA polymerase gene. How this changes impact the phage fitness and phenotype remains to be investigated.

Although most genes in the cassette had only very distant or no affiliations to genes in the databases, both the RHS-repeat and CdtB toxins had many close hits in the NCBI’s nr database (see Materials and Methods). In both cases, they formed a well supported monophyletic group (fig. 2). As aforementioned, the RHS-repeat toxin is, to the best of our knowledge, restricted to APSE-3 types. On the contrary, the CdtB protein is actually present in several APSE phages, namely APSE-7 and APSE-2. The CdtB proteins encoded in the two different APSE types do not form distinct sub-clades, rather the proteins of APSE-2 are nested within CdtB from APSE-7 (fig. 2*B*). Due to previous evidence for intragenic recombination (Degnan and Moran, 2007), we ran a recombination test using PhiPack (Bruen *et al.*, 2006) and found no significant evidence for recombination (Φ_*ω*_ p-value=7.02e-02). However, NSS- and Max χ^2^-tests did found significant recombination among CdtB genes (p-value=4.00e-03 and p-value=2.00e-03, respectively), and thus, the relationships among APSE-encoded CdtB proteins should be interpreted with caution.

**Figure 2.**
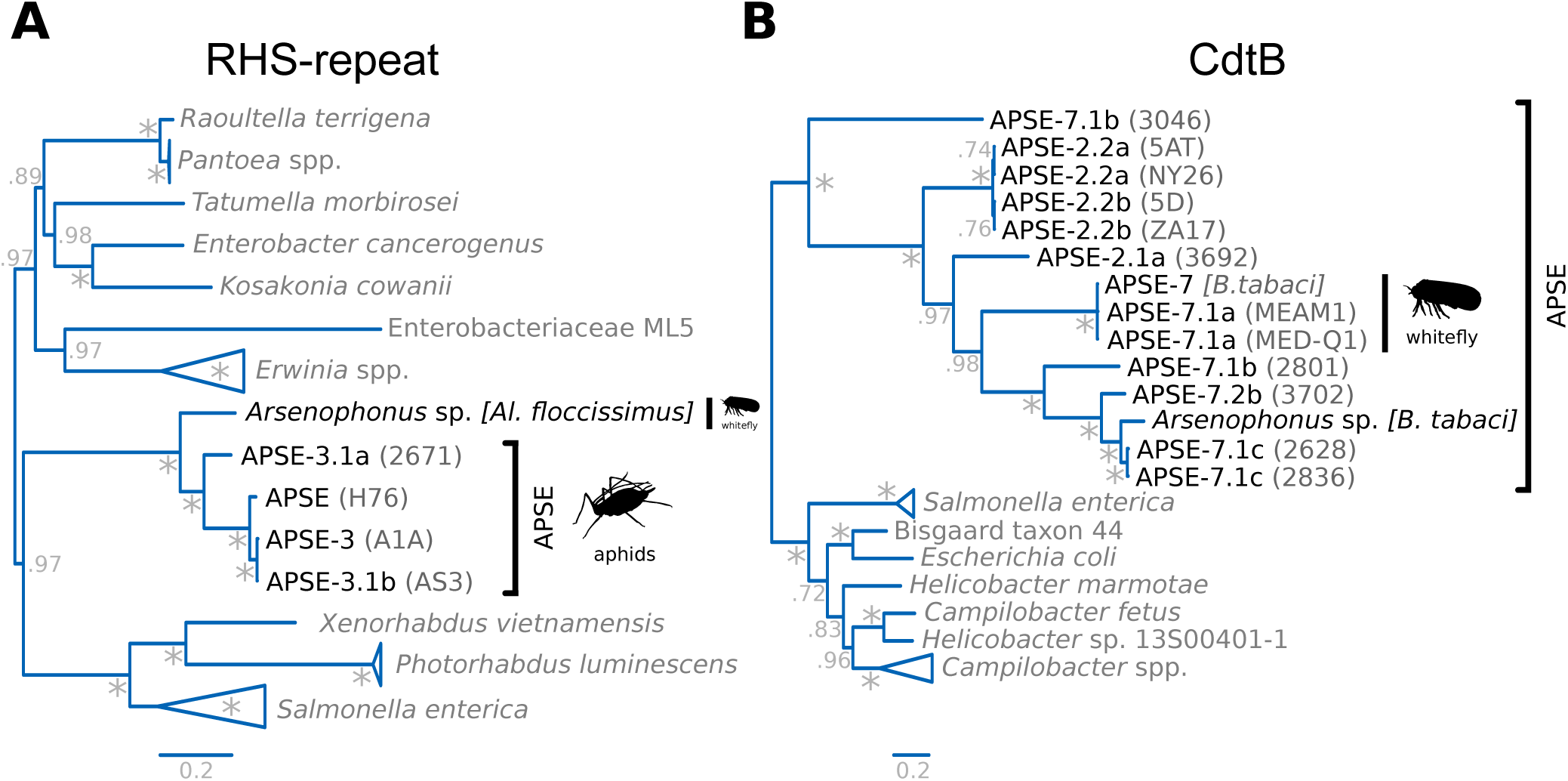
APSE toxin gene phylogenies. Bayesian phylogeny of **(A)** RHS repeat toxins and **(B)** CdtB proteins. Numbers at nodes represent Bayesian posterior probabilities. ”*”= 1. Phylogenies were midpoint rooted.

In both cases, these proteins were closely related to CdtB encoded by *Arsenophonus* spp. hosted by two whitefly species: *Aleurodicus floccissimus* and *B. tabaci*. Additionally, another four proteins in the toxin cassettes had high similarity and a close phylogenetic relation to proteins in *Arsenophonus* spp (orthogroups APSEtp_002, APSEtp_007, APSEtp_013, and APSEtp_014). Previous analyses have identified APSE-like DNA polymerase genes in *Arsenophonus* symbionts of whiteflies, aphids, louse flies, bat flies, and psyllids (Duron, 2014; Hansen *et al.*, 2007). The diagnostic PCRs used in these studies targeted the so-called P45 (DNA polymerase) and P3 (Virulence-associated protein E family protein). Analysis of three available high-quality *Arsenophonus* genomes (*Arsenophonus nasoniae* strain FIN [**CP038613.1CP038621.1**], *Arsenophonus* endosymbiont of *Nilaparvata lugens* strain Hangzhou [**JRLH01000001.1**], and *Arsenophonus* endosymbiont of *Al. floccissimus* [**OUND01000000.1**]), revealed that these reported hits actually corresponded to prophage regions in their genomes (**PHASTER** annotations available in https://doi.org/10.5281/zenodo.3764739, last accessed April 24, 2020). On closer inspection we found that only one region in the chromosome of *Ar. nasoniae* strain FIN contained both of these proteins. This region, as did others, preserved high sequence identity to one or a few APSE proteins. However, the regions were not in fact APSE phages: i.e. they consistently showed a different gene order, insertion of proteins not characteristic of fully sequenced APSE phages, and/or pseudogenenised proteins that are otherwise conserved across APSE analysed in this study (including the DNA polymerase). While there are currently no full genome sequences for the strains analysed by Hansen *et al.* 2007 nor Duron 2014, this genomic scan, along with the presence of putative toxins in *Arsenophonus* spp. closely related to those present in APSE, strongly suggests that *Arsenophonus* bacteria have indeed been historically infected by close relatives of the *Hamiltonella* APSE phages, and that these phages have left an imprint in the bacterial genomes. The retention of RHS-repeat- and CdtB-APSE-like toxin genes in at least two different strains of *Arsenophonus* suggests that these proteins could in fact endow the *Arsenophonus* symbiont with a protective phenotype. This could in turn provide a positive “APSE-free” fitness effect of carrying the endosymbiont when confronted with the environmental pressure of parasitoid infection. So far *Arsenophonus*, while quite widespread and very diversified in insects (Nováková *et al.*, 2009), has not been credited with conferring protection against parasitoids of its insect hosts.

### Phylogenetic history of *Hamiltonella* and APSE

Phylogenetic analyses supported previous observations (Degnan and Moran, 2007) that closely-related *Hamiltonella* strains were harboured by distantly related aphid species (i.e. aphids belonging to distinct subfamilies) and hosted different APSE types (fig. 3*A*). A noteworthy case is that of *Hamiltonella* strains ZA17 and 5D: they belong to two different clades but host nearly identical APSE phages. Conversely, the closely related *Hamiltonella* strain 5D and MI47 host rather different APSE types. APSE-7 is notably present in both *Hamiltonella* clade C and in the distantly related lineages of this symbiont hosted by the whitefly *B. tabaci*. This provides strong evidence and supports previous findings, based on single-gene phylogenies (Degnan and Moran, 2007, 2008), of horizontal transfer of these viral entities across *Hamiltonella* strains and lineages. The horizontal transfer of phages across lineages of endosymbiotic bacteria has been suggested for *Wolbachia*, where horizontal transfers of the WO phage seem to occur across both related and divergent *Wolbachia* (Bordenstein, 2004; Masui *et al.*, 2000). This horizontal transfer could be facilitated by the coexistence of two divergent *Hamiltonella* lineages within the same aphid host. Indeed, multiple symbiont species and strains of the same species can coexist within the same population of an aphid species (Haynes *et al.*, 2003; Meseguer *et al.*, 2017; Russell *et al.*, 2013; Sandström *et al.*, 2001; Tsuchida *et al.*, 2002), offering an opportunity for a co-infection and transfer or recombination of their phages. Through the reconstruction of a phylogenetic network using the concatenated single-copy shared proteins of APSE, we observe evidence for recombination across APSE-types (fig. 3*B*). This is a feature that has previously been observed for these phages (Degnan and Moran, 2007) and a general feature of phage genomes (reviewed in Dion *et al.*, 2020). In fact, only 18 out of 30 single-copy shared orthologous groups of proteins showed no significant evidence of intragenic recombination based on a Φ_*ω*_ test (supplementary table S4, Supplementary Material online). From these 18, nine showed significant evidence of recombination as judged by two additional tests implemented in **PhiPack**: NSS and Max χ^2^. A concatenated nucleotide alignment of the nine putative non-recombinant genes revealed significant intergenic recombination (Φ_*ω*_ p-value=0.00e+00). These results suggest genome-wide recombination for APSE phages, which is expected to complicate the inference of phylogenetic relationships. Regardless, Bayesian phylogenies were reconstructed for the three sets of single-copy shared proteins and can be found in supplementary fig. S3. Given the rampant recombination of their genomes, it is therefore hard to conclude on the phylogenetic history of these viruses.

**Figure 3.**
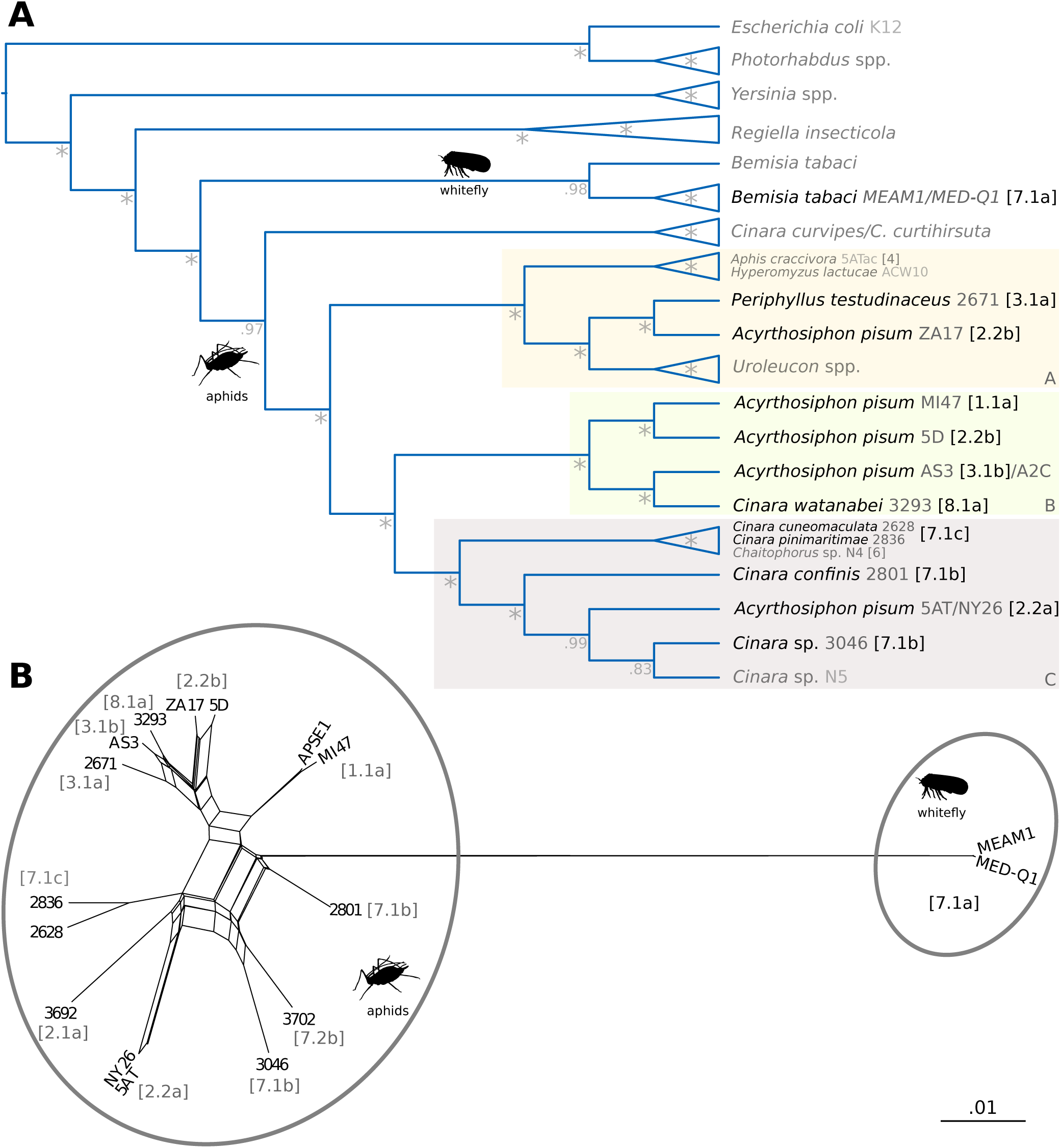
Phylogenetic relations of *Hamiltonella* and APSE phages. **(A)** Dendrogram of phylogenetic relationships among *Hamiltonella* endosymbionts based on single-copy shared proteins. Numbers at nodes represent Bayesian posterior probabilities. ”*”= 1. **(B)** Phylogenetic network of single-copy shared proteins as calculated by **SplitsTree**.

One common feature of all phylogenetic analyses was that the APSE phages infecting *B.tabaci*-associated *Hamiltonella* strains were consistently highly divergent from those of aphid-associated ones. Additionally, the **SplitsTree** network reconstruction suggests little recombination (relative to that observed among aphid-associated APSE) between these two groups of APSE infecting different hosts (aphids and *B. tabaci*). This is congruent with the *Hamiltonella* symbiont phylogeny which positions those infecting the whitefly *B. tabaci* as a separate sister clade to that made up of all currently available aphid-hosted ones (fig. 3*A*). Previous experimental studies of *Hamiltonella* infections in *Sitobion avenae* aphids have shown that strains of this symbiont establish infections more easily when transferred from the same host species (Łukasik, 2015). Additionally, these infections were found to be more stable. The same study also found that infection success was higher when the transferred symbiont strain was more closely related to the one that was originally present in the host. All this strongly suggests that these two lineages infecting different host groups can follow separate evolutionary histories, possibly driven by both ecological factors and other barriers limiting the horizontal transfer of the two distinct symbiont lineages.

### Conclusion

Facultative symbionts in insects can confer conditional fitness benefits to their hosts (Cass *et al.*, 2016; Doremus *et al.*, 2018; Polin *et al.*, 2015), and *Hamiltonella* and its APSE-conferred defence against parasitoid wasps is one of the best studied cases (Oliver and Higashi, 2019). The results presented in this study support previous studies reporting a variable toxin cassette across APSE as well as intra-genic recombination. Additionally, the whole-genome sequencing, annotation, and phylogenetic analyses of three new APSE types conducted here has revealed important features of these protective entities: (***i***) their genomes are highly conserved in terms of gene content with most of the variation localised to the toxin cassette; (***ii***) alterations can occur within a cassette, with pseudogenisation events affecting the putative toxin genes (i.e. APSE-2.2a) or IS-mediated insertion of novel genes; (***iii***) APSE harboured by whitefly-associated *Hamiltonella* seem to have undergone a separate evolutionary history, in line with their hosts; and (***iv***) *Arsenophonus* spp. might have historically had association with APSE-like phages, with some of these having left an imprint in their genomes. This study provides the first genome-wide overview of the diversity of five APSE phage types hosted by *Hamiltonella* endosymbionts, improving our understanding of these phages. The features highlighted in this study will be of great value in the design and interpretation of future experiments of APSE-mediated *Hamiltonella* parasitoid defence.

## Supporting information

File S3. Percent identity matrices for conserved APSE proteins

## Acknowledgements

This work was supported by a PhD grant from the *INRAe ECOFA department* to J.R, the *Marie-Curie AgreenSkills+* fellowship programme co-funded by the *EU’s Seventh Framework Programme* (FP7-609398) to A.M.M, the *Marie Skłodowska-Curie* fellowship program *H2020-MSCA-IF-2016* (MicroPhan: 746189) to E.J, the *France Génomique* National Infrastructure, funded as part of the *Investissemnt d’Avenir* program managed by the *Agence Nationale pour la Recherche* (ANR-10-INBS-09) to C.C. This publication has been written with the support of the AgreenSkills+ fellowship programme which has received funding from the EU’s Seventh Framework Programme under grant agreement No. FP7-609398 (AgreenSkills+ contract). We are grateful to the genotoul bioinformatics platform Toulouse Midi-Pyrenees (Bioinfo Genotoul) for providing help and/or computing and/or storage resources. The authors are grateful to the CBGP-HPC computational platform. The funders had no role in study design, data collection and analysis, decision to publish, or preparation of the manuscript.

## Data Deposition

The annotated genomes have been submitted to the European Nucleotide Archive with project numbers PRJEB15504, PRJEB31183, PRJEB37342, PRJEB37357, and PRJEB37362-PRJEB37365. The low-coverage *Hamiltonella* drafts as well as assembled and re-annotated references for APSE phages coming from previously published data are available in https://doi.org/10.5281/zenodo.3764739 (last accessed April 24, 2020).

## References

Almagro Armenteros J. J, Tsirigos K. D, Sønderby C. K, Petersen T. N, Winther O, Brunak S, von Heijne G, and Nielsen H. 2019. SignalP 5.0 improves signal peptide predictions using deep neural networks. Nature Biotechnology, 37(4): 420–423. URL https://dx.doi.org/10.1038/s41587-019-0036-z.

Altschul S. 1997. Gapped BLAST and PSI-BLAST: a new generation of protein database search programs. Nucleic Acids Res, 25(17): 3389–3402. URL https://dx.doi.org/10.1093/nar/25.17.3389.

Arndt D, Grant J. R, Marcu A, Sajed T, Pon A, Liang Y, and Wishart D. S. 2016. PHASTER: a better, faster version of the PHAST phage search tool. Nucleic Acids Research, 44(W1): W16–W21. URL https://dx.doi.org/10.1093/nar/gkw387.

Asplen M. K, Bano N, Brady C, Desneux N, Hopper K. R, Malouines C, Oliver K. M, White J. A, and Heimpel G. E. 2014. Specialisation of bacterial endosymbionts that protect aphids from parasitoids. Ecological Entomology, 39(6): 736–739. URL https://dx.doi.org/10.1111/een.12153.

Bailly-Bechet M, Vergassola M, and Rocha E. 2007. Causes for the intriguing presence of tRNAs in phages. Genome Research, 17(10): 1486–1495. URL https://dx.doi.org/10.1101/gr.6649807.

Bankevich A, Nurk S, Antipov D, Gurevich A. A, Dvorkin M, Kulikov A. S, Lesin V. M, Nikolenko S. I, Pham S, Prjibelski A. D, Pyshkin A. V, Sirotkin A. V, Vyahhi N, Tesler G, Alekseyev M. A, and Pevzner P. A. 2012. SPAdes: a new genome assembly algorithm and its applications to single-cell sequencing. Journal of Computational Biology, 19(5): 455–77. URL https://dx.doi.org/10.1089/cmb.2012.0021.

Boratyn G. M, Schäffer A. A, Agarwala R, Altschul S. F, Lipman D. J, and Madden T. L. 2012. Domain enhanced lookup time accelerated BLAST. Biol Direct, 7(1): 12. URL https://dx.doi.org/10.1186/1745-6150-7-12.

Bordenstein S. R. 2004. Bacteriophage flux in endosymbionts (*Wolbachia*): infection frequency, lateral transfer, and recombination rates. Molecular Biology and Evolution, 21(10): 1981–1991. URL https://dx.doi.org/10.1093/molbev/msh211.

Bourret J, Alizon S, and Bravo I. G. 2019. COUSIN (COdon Usage Similarity INdex): A normalized measure of codon usage preferences. Genome Biology and Evolution, 11(12): 3523–3528. URL https://dx.doi.org/10.1093/gbe/evz262.

Brandt J. W, Chevignon G, Oliver K. M, and Strand M. R. 2017. Culture of an aphid heritable symbiont demonstrates its direct role in defence against parasitoids. Proceedings of the Royal Society B: Biological Sciences, 284(1866): 20171925. URL https://dx.doi.org/10.1098/rspb.2017.1925.

Bruen T. C, Philippe H, and Bryant D. 2006. A simple and robust statistical test for detecting the presence of recombination. Genetics, 172(4): 2665–2681. URL https://dx.doi.org/10.1534/genetics.105.048975.

Cass B. N, Himler A. G, Bondy E. C, Bergen J. E, Fung S. K, Kelly S. E, and Hunter M. S. 2016. Conditional fitness benefits of the Rickettsia bacterial symbiont in an insect pest. Oecologia, 180(1): 169–179. URL https://dx.doi.org/10.1007/s00442-015-3436-x.

Chan P. P, Lin B. Y, Mak A. J, and Lowe T. M. 2019. tRNAscan-SE 2.0: Improved Detection and Functional Classification of Transfer RNA Genes. bioRxiv, page 614032. URL https://www.biorxiv.org/content/10.1101/614032v1.

Charles H and Ishikawa H. 1999. Physical and Genetic Map of the Genome of *Buchnera*, the Primary Endosymbiont of the Pea Aphid *Acyrthosiphon pisum*. J Mol Evol, 48(2): 142–150. URL http://link.springer.com/10.1007/PL00006452.

Chen F, Mackey A. J, Vermunt J. K, and Roos D. S. 2007. Assessing Performance of Orthology Detection Strategies Applied to Eukaryotic Genomes. PLoS ONE, 2(4): e383. URL https://dx.doi.org/10.1371/journal.pone.0000383.

Chen X. L, Tang D. J, Jiang R. P, He Y. Q, Jiang B. L, Lu G. T, and Tang J. L. 2011. sRNA-Xcc1, an integron-encoded transposon- and plasmid-transferred trans-acting sRNA, is under the positive control of the key virulence regulators HrpG and HrpX of *Xanthomonas campestris* pathovar *campestris*. RNA Biology, 8(6): 947–953. URL https://dx.doi.org/10.4161/rna.8.6.16690.

Chevignon G, Boyd B. M, Brandt J. W, Oliver K. M, and Strand M. R. 2018. Culture-facilitated comparative genomics of the facultative symbiont *Hamiltonella defensa*. Genome Biology and Evolution, 10(3): 786–802. URL https://dx.doi.org/10.1093/gbe/evy036/4857210.

Citron M and Schuster H. 1990. The c4 repressors of bacteriophages P1 and P7 are antisense RNAs. Cell, 62(3): 591–598. URL https://dx.doi.org/10.1016/0092-8674(90)90023-8.

Darriba D, Posada D, Kozlov A. M, Stamatakis A, Morel B, and Flouri T. 2020. ModelTest-NG: A new and scalable tool for the selection of DNA and protein evolutionary models. Molecular Biology and Evolution, 37(1): 291–294. URL https://dx.doi.org/10.1093/molbev/msz189.

Degnan P. H and Moran N. A. 2007. Evolutionary genetics of a defensive facultative symbiont of insects: exchange of toxin-encoding bacteriophage. Molecular Ecology, 17(3): 916–929. URL https://dx.doi.org/10.1111/j.1365-294X.2007.03616.x.

Degnan P. H and Moran N. A. 2008. Diverse phage-encoded toxins in a protective insect endosymbiont. Applied and Environmental Microbiology, 74(21): 6782–6791. URL https://dx.doi.org/10.1128/AEM.01285-08.

Dennis A. B, Patel V, Oliver K. M, and Vorburger C. 2017. Parasitoid gene expression changes after adaptation to symbiont-protected hosts. Evolution, 71(11): 2599–2617. URL https://dx.doi.org/10.1111/evo.13333.

Dion M. B, Oechslin F, and Moineau S. 2020. Phage diversity, genomics and phylogeny. Nature Reviews Microbiology, 18(3): 125–138. URL https://dx.doi.org/10.1038/s41579-019-0311-5.

Doremus M. R and Oliver K. M. 2017. Aphid heritable symbiont exploits defensive mutualism. Applied and Environmental Microbiology, 83(8): e03276–16. URL https://dx.doi.org/10.1128/AEM.03276-16.

Doremus M. R, Smith A. H, Kim K. L, Holder A. J, Russell J. A, and Oliver K. M. 2018. Breakdown of a defensive symbiosis, but not endogenous defences, at elevated temperatures. Molecular Ecology, 27(8): 2138–2151. URL https://dx.doi.org/10.1111/mec.14399.

Duron O. 2014. *Arsenophonus* insect symbionts are commonly infected with APSE, a bacteriophage involved in protective symbiosis. FEMS Microbiology Ecology, 90(1): 184–194. URL https://dx.doi.org/10.1111/1574-6941.12381.

Edgar R. C. 2004. MUSCLE: multiple sequence alignment with high accuracy and high throughput. Nucleic Acids Research, 32(5): 1792–1797. URL https://dx.doi.org/10.1093/nar/gkh340.

Gouy M, Guindon S, and Gascuel O. 2010. SeaView Version 4: A multiplatform graphical user interface for sequence alignment and phylogenetic tree building. Molecular Biology and Evolution, 27(2): 221–224. URL https://dx.doi.org/10.1093/molbev/msp259.

Guo J, Hatt S, He K, Chen J, Francis F, and Wang Z. 2017. Nine facultative endosymbionts in aphids. A review. Journal of Asia-Pacific Entomology, 20(3): 794–801. URL https://dx.doi.org/10.1016/j.aspen.2017.03.025.

Hansen A. K, Jeong G, Paine T. D, and Stouthamer R. 2007. Frequency of secondary symbiont infection in an invasive psyllid relates to parasitism pressure on a geographic scale in California. Applied and Environmental Microbiology, 73(23): 7531–7535. URL https://dx.doi.org/10.1128/AEM.01672-07.

Haynes S, Darby A. C, Daniell T. J, Webster G, van Veen F. J. F, Godfray H, Prosser J. I, and Douglas A. E. 2003. Diversity of bacteria associated with natural aphid populations. Applied and Environmental Microbiology, 69(12): 7216–7223. URL https://dx.doi.org/10.1128/AEM.69.12.7216-7223.2003.

Hopper K. R, Kuhn K. L, Lanier K, Rhoades J. H, Oliver K. M, White J. A, Asplen M. K, and Heimpel G. E. 2018. The defensive aphid symbiont *Hamiltonella defensa* affects host quality differently for *Aphelinus glycinis* versus *Aphelinus atriplicis*. Biological Control, 116: 3–9. URL https://dx.doi.org/10.1016/j.biocontrol.2017.05.008.

Jones P, Binns D, Chang H.-Y, Fraser M, Li W, McAnulla C, McWilliam H, Maslen J, Mitchell A, Nuka G, Pesseat S, Quinn A. F, Sangrador-Vegas A, Scheremetjew M, Yong S.-Y, Lopez R, and Hunter S. 2014. InterProScan 5: genome-scale protein function classification. Bioinformatics, 30(9): 1236–1240. URL https://dx.doi.org/10.1093/bioinformatics/btu031.

Jousselin E, Clamens A.-L, Galan M, Bernard M, Maman S, Gschloessl B, Duport G, Meseguer A. S, Calevro F, and Cœur d’Acier A. 2016. Assessment of a 16S rRNA amplicon Illumina sequencing procedure for studying the microbiome of a symbiont-rich aphid genus. Mol Ecol Resour, 16(3): 628–640. URL http://doi.wiley.com/10.1111/1755-0998.12478.

Kall L, Krogh A, and Sonnhammer E. L. L. 2005. An HMM posterior decoder for sequence feature prediction that includes homology information. Bioinformatics, 21(Suppl 1): i251–i257. URL https://dx.doi.org/10.1093/bioinformatics/bti1014.

Kalvari I, Nawrocki E. P, Argasinska J, Quinones-Olvera N, Finn R. D, Bateman A, and Petrov A. I. 2018a. Non-Coding RNA analysis using the Rfam database. Current Protocols in Bioinformatics, 62(1): e51. URL https://dx.doi.org/10.1002/cpbi.51.

Kalvari I, Argasinska J, Quinones-Olvera N, Nawrocki E. P, Rivas E, Eddy S. R, Bateman A, Finn R. D, and Petrov A. I. 2018b. Rfam 13.0: shifting to a genome-centric resource for non-coding RNA families. Nucleic Acids Research, 46(D1): D335–D342. URL https://dx.doi.org/10.1093/nar/gkx1038.

Katoh K and Standley D. M. 2013. MAFFT Multiple sequence alignment software version 7: Improvements in performance and usability. Mol Biol Evol, 30(4): 772–780. URL https://dx.doi.org/10.1093/molbev/mst010.

Lanfear R, Frandsen P. B, Wright A. M, Senfeld T, and Calcott B. 2017. PartitionFinder 2: New methods for selecting partitioned models of evolution for molecular and morphological phylogenetic analyses. Molecular Biology and Evolution, 34(3): 772–773. URL https://dx.doi.org/10.1093/molbev/msw260.

Lenhart P. A and White J. A. 2017. A defensive endosymbiont fails to protect aphids against the parasitoid community present in the field. Ecological Entomology, 42(5): 680–684. URL https://dx.doi.org/10.1111/een.12419.

Leybourne D. J, Bos J. I. B, Valentine T. A, and Karley A. J. 2020. The price of protection: a defensive endosymbiont impairs nymph growth in the bird cherry-oat aphid, *Rhopalosiphum padi*. Insect Science, 27(1): 69–85. URL https://dx.doi.org/10.1111/1744-7917.12606.

Li L. 2003. OrthoMCL: Identification of Ortholog Groups for Eukaryotic Genomes. Genome Res, 13(9): 2178–2189. URL https://dx.doi.org/10.1101/gr.1224503.

Łukasik P, Guo H, van Asch M, Henry L. M, Godfray H. C. J, and Ferrari J. 2015. Horizontal transfer of facultative endosymbionts is limited by host relatedness. Evolution, 69(10): 2757–2766. URL https://dx.doi.org/10.1111/evo.12767.

Manzano-Marín A, Coeur d’acier A, Clamens A.-L, Orvain C, Cruaud C, Barbe V, and Jousselin E. 2018. A freeloader? The highly eroded yet large genome of the Serratia symbiotica symbiont of Cinara strobi. Genome Biology and Evolution, 10(9): 2178–2189. URL https://dx.doi.org/10.1093/gbe/evy173.

Manzano-Marín A, Coeur d’acier A, Clamens A.-L, Orvain C, Cruaud C, Barbe V, and Jousselin E. 2020. Serial horizontal transfer of vitamin-biosynthetic genes enables the establishment of new nutritional symbionts in aphids’ di-symbiotic systems. The ISME Journal, 14(1): 259–273. URL https://dx.doi.org/10.1038/s41396-019-0533-6.

Martinez A. J, Weldon S. R, and Oliver K. M. 2014. Effects of parasitism on aphid nutritional and protective symbioses. Molecular Ecology, 23(6): 1594–1607. URL https://dx.doi.org/10.1111/mec.12550.

Masui S, Kamoda S, Sasaki T, and Ishikawa H. 2000. Distribution and evolution of bacteriophage WO in Wolbachia, the endosymbiont causing sexual alterations in arthropods. Journal of Molecular Evolution, 51(5): 491–497. URL https://dx.doi.org/10.1007/s002390010112.

Meseguer A. S, Manzano-Marín A, Coeur d’Acier A, Clamens A.-L, Godefroid M, and Jousselin E. 2017. Buchnera has changed flatmate but the repeated replacement of co-obligate symbionts is not associated with the ecological expansions of their aphid hosts. Molecular Ecology, 26(8): 2363–2378. URL https://dx.doi.org/10.1111/mec.13910.

Milne I, Stephen G, Bayer M, Cock P. J. A, Pritchard L, Cardle L, Shaw P. D, and Marshall D. 2013. Using Tablet for visual exploration of second-generation sequencing data. Briefings in Bioinformatics, 14(2): 193–202. URL https://dx.doi.org/10.1093/bib/bbs012.

Mitchell A. L, Attwood T. K, Babbitt P. C, Blum M, Bork P, Bridge A, Brown S. D, Chang H.-Y, El-Gebali S, Fraser M. I, Gough J, Haft D. R, Huang H, Letunic I, Lopez R, Luciani A, Madeira F, Marchler-Bauer A, Mi H, Natale D. A, Necci M, Nuka G, Orengo C, Pandurangan A. P, Paysan-Lafosse T, Pesseat S, Potter S. C, Qureshi M. A, Rawlings N. D, Redaschi N, Richardson L. J, Rivoire C, Salazar G. A, Sangrador-Vegas A, Sigrist C. J. A, Sillitoe I, Sutton G. G, Thanki N, Thomas P. D, Tosatto S. C. E, Yong S.-Y, and Finn R. D. 2019. InterPro in 2019: improving coverage, classification and access to protein sequence annotations. Nucleic Acids Research, 47(D1): D351–D360. URL https://dx.doi.org/10.1093/nar/gky1100.

Moran N. A, Degnan P. H, Santos S. R, Dunbar H. E, and Ochman H. 2005. The players in a mutualistic symbiosis: Insects, bacteria, viruses, and virulence genes. Proceedings of the National Academy of Sciences, 102(47): 16919–16926. URL https://dx.doi.org/10.1073/pnas.0507029102.

Nawrocki E. P and Eddy S. R. 2013. Infernal 1.1: 100-fold faster RNA homology searches. Bioinformatics, 29(22): 2933–2935. URL https://dx.doi.org/10.1093/bioinformatics/btt509.

Nováková E, Hypša V, and Moran N. A. 2009. *Arsenophonus*, an emerging clade of intracellular symbionts with a broad host distribution. BMC Microbiology, 9(1): 143. URL https://dx.doi.org/10.1186/1471-2180-9-143.

Okonechnikov K, Golosova O, and Fursov M. 2012. Unipro UGENE: a unified bioinformatics toolkit. Bioinformatics, 28(8): 1166–1167. URL https://dx.doi.org/10.1093/bioinformatics/bts091.

Oliver K. M and Higashi C. H. 2019. Variations on a protective theme: *Hamiltonella defensa* infections in aphids variably impact parasitoid success. Current Opinion in Insect Science, 32: 1–7. URL https://dx.doi.org/10.1016/j.cois.2018.08.009.

Oliver K. M, Campos J, Moran N. A, and Hunter M. S. 2008. Population dynamics of defensive symbionts in aphids. Proceedings of the Royal Society B: Biological Sciences, 275(1632): 293–299. URL https://dx.doi.org/10.1098/rspb.2007.1192.

Oliver K. M, Degnan P. H, Hunter M. S, and Moran N. A. 2009. Bacteriophages encode factors required for protection in a symbiotic mutualism. Science, 325(5943): 992–994. URL https://dx.doi.org/10.1126/science.1174463.

Oliver K. M, Degnan P. H, Burke G. R, and Moran N. A. 2010. Facultative symbionts in aphids and the horizontal transfer of ecologically important traits. Annual Review of Entomology, 55(1): 247–266. URL https://dx.doi.org/10.1146/annurev-ento-112408-085305.

Padalon-Brauch G, Hershberg R, Elgrably-Weiss M, Baruch K, Rosenshine I, Margalit H, and Altuvia S. 2008. Small RNAs encoded within genetic islands of *Salmonella typhimurium* show host-induced expression and role in virulence. Nucleic Acids Research, 36(6): 1913–1927. URL https://dx.doi.org/10.1093/nar/gkn050.

Patel V, Chevignon G, Manzano-Marín A, Brandt J. W, Strand M. R, Russell J. A, and Oliver K. M. 2019. Cultivation-assisted genome of *Candidatus* Fukatsuia symbiotica; the enigmatic ‘X-type’ symbiont of aphids. Genome Biology and Evolution, 11(12): 3510–3522. URL https://dx.doi.org/10.1093/gbe/evz252.

Polin S, Le Gallic J.-F, Simon J.-C, Tsuchida T, and Outreman Y. 2015. Conditional reduction of predation risk associated with a facultative symbiont in an insect. PLOS ONE, 10(11): e0143728. URL https://dx.doi.org/10.1371/journal.pone.0143728.

Rice P, Longden I, and Bleasby A. 2000. EMBOSS: The European molecular biology open software suite. Trends in Genetics, 16(6): 276–277. URL https://dx.doi.org/10.1016/S0168-9525(00)02024-2.

Ronquist F, Teslenko M, van der Mark P, Ayres D. L, Darling A, Hohna S, Larget B, Liu L, Suchard M. A, and Huelsenbeck J. P. 2012. MrBayes 3.2: Efficient Bayesian Phylogenetic Inference and Model Choice Across a Large Model Space. Syst Biol, 61(3): 539–542. URL https://dx.doi.org/10.1093/sysbio/sys029.

Russell J. A, Weldon S, Smith A. H, Kim K. L, Hu Y, Łukasik P, Doll S, Anastopoulos I, Novin M, and Oliver K. M. 2013. Uncovering symbiont-driven genetic diversity across North American pea aphids. Molecular Ecology, 22(7): 2045–2059. URL https://dx.doi.org/10.1111/mec.12211.

Sandström J. P, Russell J. A, White J. P, and Moran N. A. 2001. Independent origins and horizontal transfer of bacterial symbionts of aphids. Molecular Ecology, 10(1): 217–228. URL https://dx.doi.org/10.1046/j.1365-294X.2001.01189.x.

Schmieder R and Edwards R. 2011. Quality control and preprocessing of metagenomic datasets. Bioinformatics, 27(6): 863–864. URL https://dx.doi.org/10.1093/bioinformatics/btr026.

Seemann T. 2014. Prokka: rapid prokaryotic genome annotation. Bioinformatics, 30(14): 2068–2069. URL https://dx.doi.org/10.1093/bioinformatics/btu153.

Talavera G and Castresana J. 2007. Improvement of phylogenies after removing divergent and ambiguously aligned blocks from protein sequence alignments. Syst Biol, 56(4): 564–577. URL https://dx.doi.org/10.1080/10635150701472164.

Touchon M, Moura de Sousa J. A, and Rocha E. P. 2017. Embracing the enemy: the diversification of microbial gene repertoires by phage-mediated horizontal gene transfer. Current Opinion in Microbiology, 38: 66–73. URL https://dx.doi.org/10.1016/j.mib.2017.04.010.

Tsuchida T, Koga R, Shibao H, Matsumoto T, and Fukatsu T. 2002. Diversity and geographic distribution of secondary endosymbiotic bacteria in natural populations of the pea aphid, *Acyrthosiphon pisum*. Molecular Ecology, 11(10): 2123–2135. URL https://dx.doi.org/10.1046/j.1365-294X.2002.01606.x.

van der Wilk F, Dullemans A. M, Verbeek M, and van den Heuvel J. F. 1999. Isolation and characterization of APSE-1, a bacteriophage infecting the secondary endosymbiont of *Acyrthosiphon pisum*. Virology, 262(1): 104–113. URL https://dx.doi.org/10.1006/viro.1999.9902.

Weldon S. R, Strand M. R, and Oliver K. M. 2013. Phage loss and the breakdown of a defensive symbiosis in aphids. Proceedings of the Royal Society B: Biological Sciences, 280(1751): 20122103. URL https://dx.doi.org/10.1098/rspb.2012.2103.

Worning P, Jensen L. J, Hallin P. F, Stærfeldt H.-H, and Ussery D. W. 2006. Origin of replication in circular prokaryotic chromosomes. Environmental Microbiology, 8(2): 353–361. URL https://dx.doi.org/10.1111/j.1462-2920.2005.00917.x.

Zemskov E. A, Kang W, and Maeda S. 2000. Evidence for nucleic acid binding ability and nucleosome association of *Bombyx mori* nucleopolyhedrovirus BRO proteins. Journal of Virology, 74(15): 6784–6789. URL https://dx.doi.org/10.1128/JVI.74.15.6784-6789.2000.

Zhou Y, Liang Y, Lynch K. H, Dennis J. J, and Wishart D. S. 2011. PHAST: A fast phage search tool. Nucleic Acids Research, 39(suppl): W347–W352. URL https://dx.doi.org/10.1093/nar/gkr485.

