## Supplementary material for "The protector within: Comparative genomics of APSE phages across aphids reveals rampant recombination and diverse toxin arsenals": File S2. Supplementary Figures S1-S3

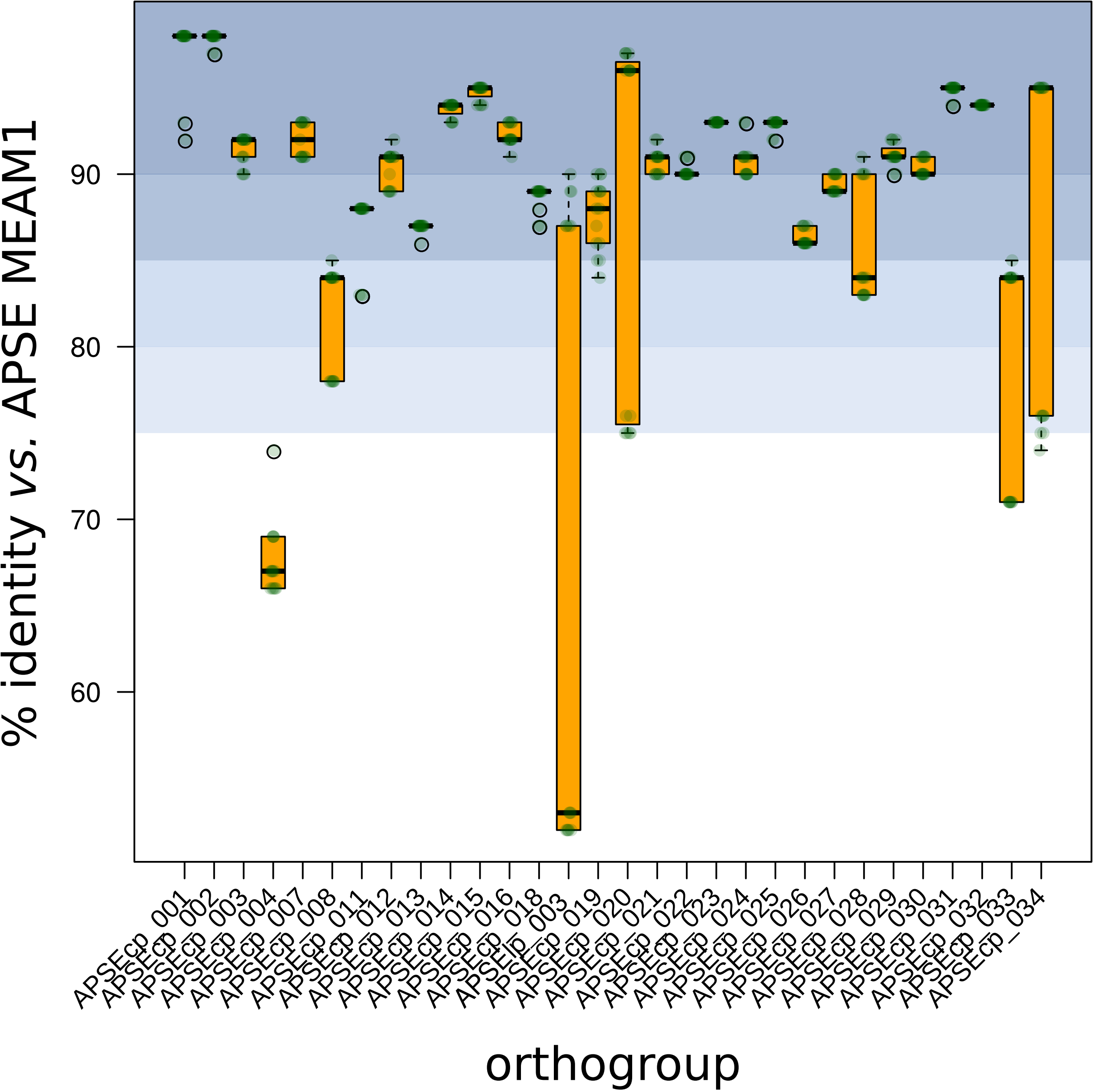


**Fig. S1. Identity of conserved proteins across APSE *vs.* MEAM1.** Example of identity across single-copy shared proteins across APSE phages *vs.* MEAM1 as caclulated in ClustalX. A file showing identity tables across all 30 proteins can be found in file S3.

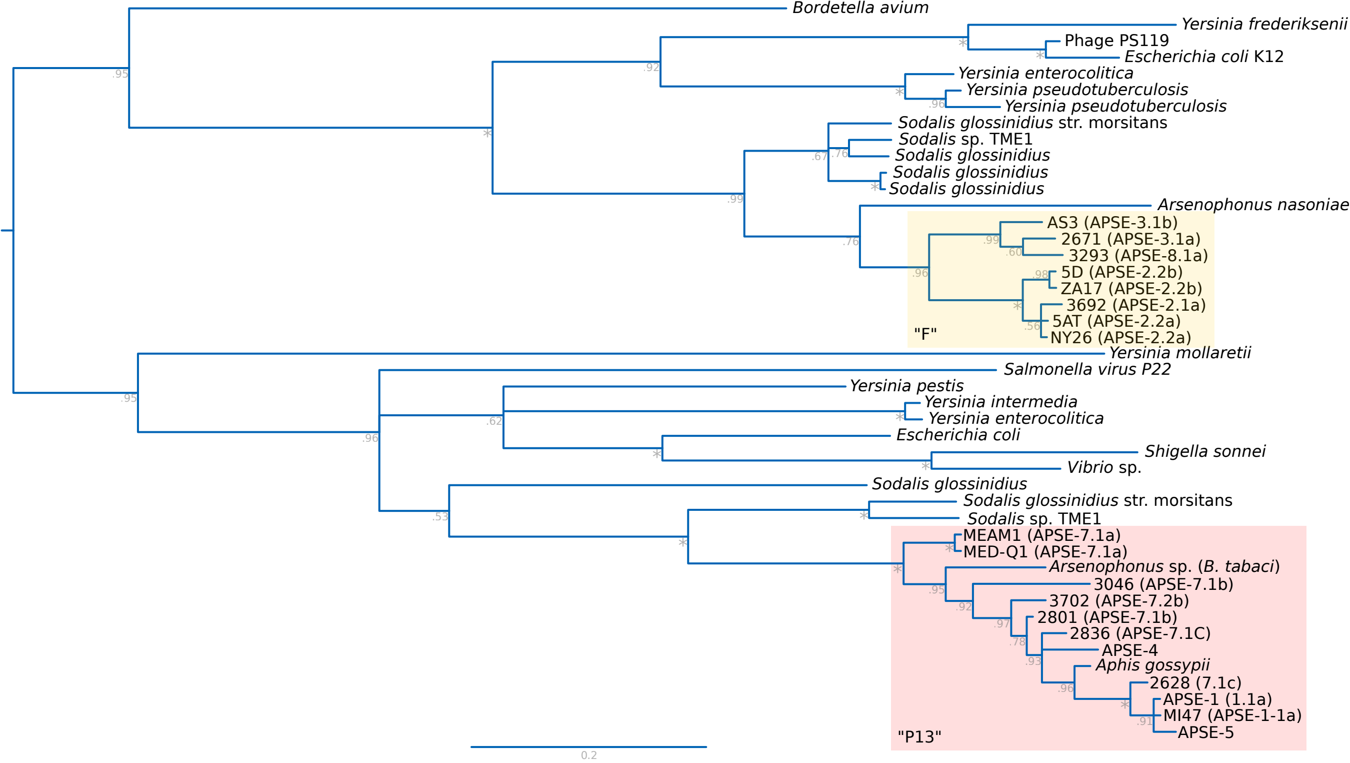


**Fig. S2. Bayesian phylogeny of lyzozyme genes from APSE phages.** Amino acid Bayesian phylogeny reconstructed in MrBayes v3.2.7 (JTT+I+G4; 300,000 generations) highlighting the two lyzozyme phylogroups: the so-called "F" (yellow) and "P13" (red). Phylogeny was midpoint rooted.

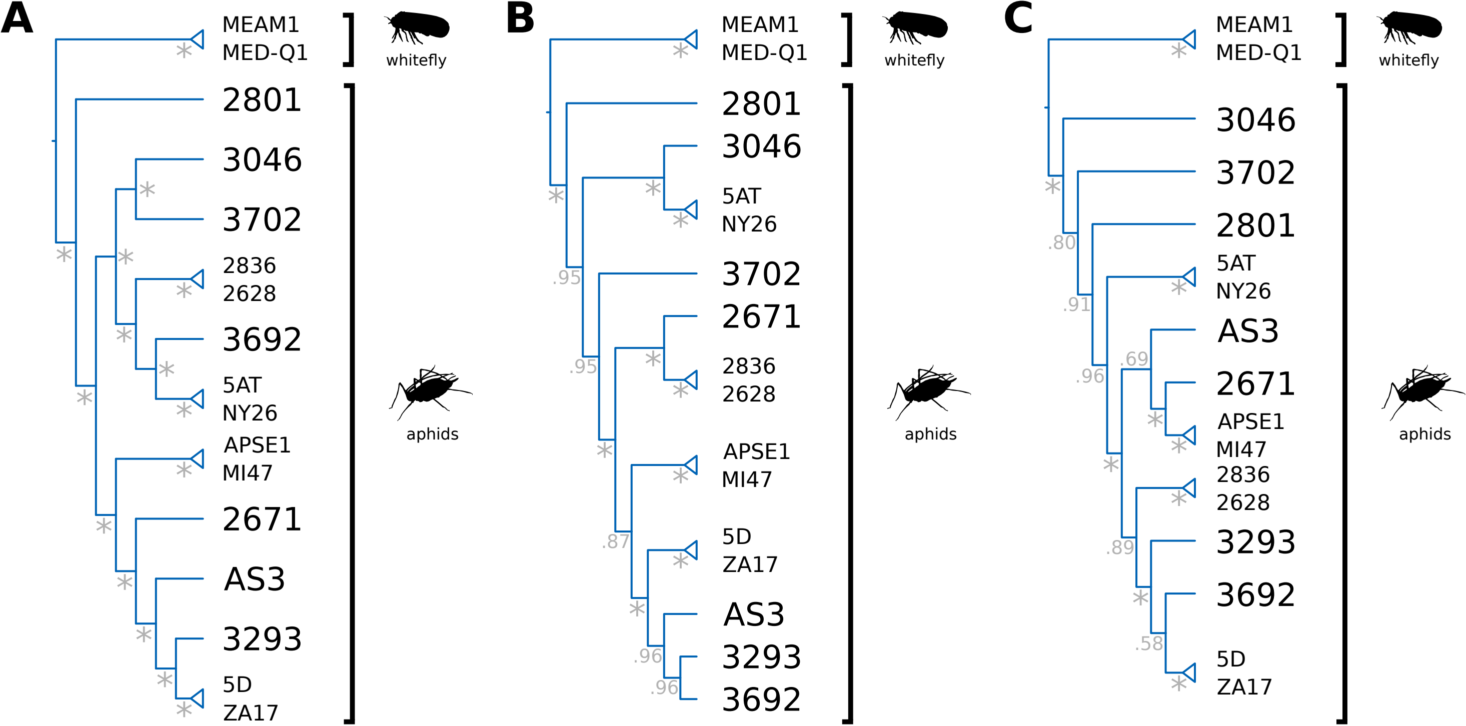


**Fig. S3. Bayesian phylogenies of single-copy shared proteins of APSE phages.** Amino acid Bayesian phylogenies reconstructed in MrBayes v3.2.7 for (A) all 30 single-copy shared orthologous groups of proteins, (B) all 18 non-recombinant ones based on $\Phi_{\omega}$, and (C) all 9 showing additionally no statistically significant recobmination in two other tests, NSS and Max $\chi^2$. Asterisks at nodes represent a posterior probability of 1. APSE strain name is shown at tips.
